# The genomic formation of Tanka people, an isolated “Gypsies in water” in the coastal region of Southeast China

**DOI:** 10.1101/2021.07.18.452867

**Authors:** Guanglin He, Yunhe Zhang, Lan-Hai Wei, Mengge Wang, Xiaomin Yang, Jianxin Guo, Rong Hu, Chuan-Chao Wang, Xian-Qing Zhang

**Affiliations:** Department of Anthropology and Ethnology, Institute of Anthropology, National Institute for Data Science in Health and Medicine, State Key Laboratory of Cellular Stress Biology, School of Life Sciences, State Key Laboratory of Marine Environmental Science, Xiamen University, Xiamen, 361005, China; School of Humanities, Nanyang Technological University, Nanyang Avenue, 639798, Singapore; School of Public Administration, Zhejiang Gongshang University, Hangzhou, 310018, China; Guangzhou Forensic Science Institute, Guangzhou, 510080, China; Faculty of Forensic Medicine, Zhongshan School of Medicine, Sun Yat-sen University, Guangzhou, 510080, China; School of Basic Medical Sciences, Zhejiang University School of Medicine, Hangzhou, 310000, China; Institute of Asian Civilizations, Zhejiang University, Hangzhou, 310000, China; B&R International Joint Laboratory for Eurasian Anthropology, Fudan University, Shanghai, 200438, China

**Keywords:** Tanka people, genetic history, admixture events, population genomics, molecular anthropology

## Abstract

**Objectives:** Three different hypotheses proposed via the controversial evidence from cultural, anthropological and uniparental genetic analysis respectively stated that Tanka people probably originated from Han Chinese, ancient Baiyue tribe, or the admixture of them. Therefore, the genetic origin and admixture history of the Tanka people, an isolated “Gypsies in water” in the coastal region of Southeast China, are needed to be genetically clarified.

**Materials and methods:** To elucidate the genetic origin of the Southeast Tanka people and explore their genetic relationship with surrounding indigenous Tai-Kadai and Austronesian people and Neolithic-to-historic ancients from the Yellow River Basin (YRB) and Fujian, we conducted a large-scale population genomic study among 1498 modern and ancient Eurasians, in which 73 Tanka and 4 Han people were first reported here. Both allele-shared and haplotype-based statistical methods were used here, including PCA, ADMIXTURE, *f*-statistics, ALDER, *qpGraph-/TreeMix and qpAdm/qpWave*, ChromoPainter and FineSTRUCTURE

**Results:** We found a specific genetic cline in PCA plots and detected the Tanka-specific homogeneous ancestry in model-based ADMIXTURE results, suggesting differentiated demographic history between Tanka and surrounding Hans. Formal tests based on sharing allele patterns showed a close relationship between Tanka people and Han Chinese, but the Tanka population harbored more southern East Asian ancestry related to Austronesian and Tai-Kadai people compared with southern Hans. Besides, the reconstructed differentiated demographic history revealed that southern Xinshizhou Tanka harbored more ancestry related to the Tai-Kadai or coastal ancient Neolithic to Bronze Age East Asians compared with northern Shacheng Tanka. The *qpGraph-/TreeMix*-based phylogenetic framework, *qpAdm/qpWave*-based admixture modeling and FineSTRUCTURE-based dendrogram among ancient northern and southern East Asians further demonstrated that the primary ancestry of modern Tanka derived from ancient millet farmers in the YRB with additional admixture from multiple southern East Asian sources.

**Discussion:** Sharing ancestry estimated from the *f*-statistics and sharing haplotypic landscape inferred from the ChromoPainter and FineSTRUCTURE showed that Southeast Tanka people not only had a close genetic relationship with both Northern Hans and YRB millet farmers but also possessed more southern East Asian ancestry related to Austronesian, Tai-Kadai and Hmong-Mien people. Our genomic data and fitted admixture models supported modern Tanka originated from ancient North China and obtained additional gene flow from ancient southern East Asians in the processes of southward migrations.

## INTRODUCTION

The overwhelming southward population dispersal from the central plain around the Yellow River Basin (**YRB**) of North-East Asia in the past two thousand years resulted in the formation of Han populations in southern China (Guang-Lin He et al., 2021; G. He et al., 2020; J. Sun et al., 2021). Meanwhile, it is also generally accepted that this process was accompanied by a large scale of integration with the southern aborigines (Guang-Lin He et al., 2021; G. He et al., 2020; J. Sun et al., 2021). There are many different local clans of Han populations in different geographic regions of southern China. Some of these people have very special cultural traditions so that scholars speculate that they may be descendants of remote sub-branches of the ancestor of Han populations or have a high proportion of genetic elements from the southern indigenous people (S. Liu et al., 2018; J. Sun et al., 2021; Wen et al., 2004), such minority groups including Chuanqing, Gejia, Dongjia, Xijia, etc. (Lu et al., 2020; Jin Sun et al., 2020). The Tanka people, also called the “Gypsies in water” in the coastal region of southeastern China, were also considered as a special clan of Han populations. Further population genetic study focused on the origin of these minority populations will assist in exploring the detailed genetic history to better understand the population demographic history in South China, as well as better reconstructing the formation process of Han populations in the past two thousand years.

There are long debates about the ancient origin of Tanka people, mainly focused on three hypotheses of the northern Han Chinese origin, southern indigenous Baiyue origin, and the mixture of them (Luo et al., 2020). Modern Tanka people are widely scattered in the coastal region of Southeast China, ranging from Zhejiang province to Guangxi province. Historically documented evidence showed that Tanka people were the decedents of the ancient Baiyue tribe and subsequently continuously stimulated by the southward migrated Han Chinese populations from the central plain, which was further confirmed via physical anthropological features. Supporting evidence of the ancient Baiyue origin of the Tanka people was also provided in the perspective of uniparental genetic legacy (Luo et al., 2020). Luo et al. genotyped both maternally and paternally informative SNPs in Fujian Tanka people and explored their patrilineal and matrilineal genetic history (Luo et al., 2020). This uniparental genetic legacy investigation found two predominant Y-chromosome lineages of O1a1a-P203 and O1b1a1a-M95 and three maternal founding lineages of F2a, M7c1 and F1a1. Patterns of genetic affinity via clustering technique of principal component analysis revealed the contentious relationships: a close relationship with Tai-Kadai speakers based on paternal genetic variations, but an affinity with Han Chinese based on the maternal genetic variations. Other analyses of divergence times based on Tanka-specific haplotypes suggested their ancient indigenous origin approximately 1000 years ago, which is not consistent with its admixture model with two sources related to the southward Han and southern indigenous people. However, the genetic legacy from the autosomal perspectives and their ancient relationship with modern and ancient East Asians keeps in its infancy.

Advances in ancient DNA studies in East Asia also provided new insights into the formation of modern and ancient East Asian populations. Wang et al. recently reconstructed one deep evolutionary framework and found that the deep paleolithic coastal migration route dispersed one deeply diverged lineages related to South Asian Onge Hunter-Gatherer people, which also widely distributed in modern and ancient Tibetans, Jomon and southeastern coastal Hanben people with a variable proportion (C. C. Wang et al., 2021). Further three Holocene expansions from Amur River Basin, YRB, and Yangtze River Basin dispersed East Asia’s language, farming and people, which reshaped the patterns of the modern mosaic genetic landscape of East Asia (Ning et al., 2020; C. C. Wang et al., 2021; Yang et al., 2020). Ancient DNA research further demonstrated significant southward migration of millet farmers from YRB to South China, as well as southward migration of southern Chinese agriculturists to the Island and Mainland of Southeast Asia, which further complicated the following molecular patterns of the population genomic diversity in ancient southern Chinese indigenous people and Southeast Asians (Larena et al., 2021; Lipson et al., 2018; McColl et al., 2018; Yang et al., 2020). These genetic legacy investigations provided better proxies or surrogates of the ancient sources for modeling the formation of modern East Asians.

Thus, we obtained the genome-wide SNPs data from the southeastern region of China and merged them with all available modern and ancient East Asians (Guanglin He et al., 2020; Lipson et al., 2018; D. Liu et al., 2020; McColl et al., 2018; Ning et al., 2020; C. C. Wang et al., 2021; Yang et al., 2020) to perform population genomic studies focused on the exploration of the genetic origin of Tanka people and their interaction with modern and ancient surrounding populations. We identified one unique ancestry composition in the Tanka people, which can be modeled as the admixture result of the southern sources related to the coastal Austronesian-speaking Ami/ancient Hanben, and northern sources related to Northern Han or Neolithic YRB farmers. Clustering results showed their relatively isolated position in PCA, and unique admixture signatures in ADMIXTURE, and a closer relationship with southern indigenes than with southern Han Chinese populations. Besides, *f*_*4*_-statistics, *qpAdm, qpGraph* and FineSTRUCTURE results further revealed that Tanka not only harbored a close relationship with modern southern Han Chinese but also had a close connection with ancient Yellow River Basin farmers, as well as southern Late Neolithic to Iron Age Hanben, which can be modeled as the result of major northern East Asian ancestry related to millet farmers and minor southern East Asian ancestry related to Hanben. Our genomic evidence supported the ‘admixture hypothesis’ with one source related to the northern East Asians and the other related to the southern indigenes.

## MATERIALS AND METHODS

### Sample collections, genotyping and quality control

We collected 77 saliva samples from unrelated healthy individuals from three populations in Fujian province, southeastern China. We performed this study strictly followed by the regulations of the Human Genetic Resources Administration of China (HGRAC) and the recommendations of the Helsinki Declaration of 2000 (Association, 2001). The research protocol has been approved via the Medical Ethics Committee of Xiamen University (XDYX201909). The informed consent was signed before sample collections. We used the DP-318 Kit (Tiangen Biotechnology, Beijing) to isolate the genomic DNA based on the manufactures’ instructions. Genotype data of approximately 700K genome-wide SNPs were sequenced via the Illumina Infinium^®^ Global Screening Array (GSA). Missing SNPs and missing individuals were checked using the Plink 1.9 (Purcell et al., 2007) with the two parameters (mind: 0.01 and geno: 0.01). We then merged the quality-controlled data with previously published modern and ancient Eurasian populations included in the Human Origin dataset or 1240K dataset publicly shared from Reich Lab and other published genetic studies (Chen et al., 2021; Guanglin He et al., 2020; He et al., 2021; Lipson et al., 2018; D. Liu et al., 2020; Y. Liu et al., 2021; Lu et al., 2020; McColl et al., 2018; Ning et al., 2020; C. C. Wang et al., 2021; Q. Wang et al., 2020; Yang et al., 2020; Yao et al., 2021).

### Sharing allele-based analyses from PCA, ADMIXTURE, *f*-statistics and TreeMix

Plink 1.9 (Purcell et al., 2007) was used to prune a dataset with strong linkage disequilibrium (--indep-pairwise 200 25 0.4) and calculated the pairwise genetic distance of Fst index to evaluate the genetic similarities between three Fujian populations and other eastern Eurasians. Principal component analysis (PCA) among eastern modern and ancient Eurasians or their subsets was performed using smartpca package in the EIGENSOFT (Patterson et al., 2012) with two parameters (lsqproject: YES and numoutlieriter: 0). Unsupervised model-based ADMIXTURE analyses were carried out using ADMIXTURE 1.3.0 (Alexander, Novembre, & Lange, 2009) and pruned datasets with the predefined ancestral populations ranging from two to twenty.

All formal tests of allele-shared analysis using different packages in ADMIXTOOLS (Patterson et al., 2012). The genetic affinity between the studied Fujian populations and other Eurasian references was measured via outgroup-*f*_*3*_-statistics in the form *f*_*3*_(Eurasian modern and ancients, Fujian populations; Mbuti) using the *qp3pop* package. Admixture signals were explored via admixture-*f*_*3*_(Eurasian source1, Eurasian source2; Fujian populations). Symmetrical-*f*_*4*_-statistics in the form of *f*_*4*_(Eurasian reference population1, Eurasian reference population2; Fujian populations, Mbuti) and *f*_*4*_(Predefined ancestral source proxy, Fujian populations, Eurasian reference population, Mbuti) were calculated *qpDstat* package in ADMIXTOOLS (Patterson et al., 2012) with *f*_*4*_Mode (*f*_*4*_: YES). Population splits and gene flow events among southern modern and ancient East Asians were constructed via TreeMix (Pickrell & Pritchard, 2012) and further explored via the *qpGraph* (Patterson et al., 2012) with more complex models based on different Paleolithic, Neolithic and modern genetic variations. We also used *qpWave/qpAdm* packages in ADMIXTOOLS (Patterson et al., 2012) to evaluate the admixture proportion in our predefined three-way admixture models with northern East Asian ancients from the YRB, southern coastal East Asians from Fujian and inland East Asians from Vietnam as three source proxies. Dates of admixture events were estimated via Admixture-induced Linkage Disequilibrium for Evolutionary Relationships (ALDER) (Loh et al., 2013) based on the fitted decay rate of linkage disequilibrium. Haplogroups of Y-chromosome and mitochondria DNA were assigned via the in-house scripts.

### Sharing-haplotype-based IBD and finer-scale population dendrograms inferred from FineSTRUCTURE

We phased genome-wide dense SNP data using ShapeIT and then calculated shared Identity by Descent (IBD) between Fujian Tanka and other reference populations using the Refined IBD(Browning & Browning, 2011). We painted the Tanka’s genomes using all our predefined donor chromosomes via the Chromopaintor2 and explored the finer-scale genetic structure based on the coancestry matrix using the FineSTRUCTURE v4(Lawson, Hellenthal, Myers, & Falush, 2012). And finally, we identified the ancestry sources and dated corresponding admixture models using GLOBETROTTER(Hellenthal et al., 2014).

## RESULTS AND DISCUSSION

### Overview of genetic structure and general population relationship

We newly generated genome-wide single nucleotide polymorphism (SNP) data approximately 700,000 genetic markers in 77 southern Chinese individuals, including 73 officially ethnically unrecognized Tanka people from Shacheng (34) and Xinshizhou (39) and four Han Chinese individuals from Fujian province. We merged the new-obtained data with 997 modern eastern Eurasian from 105 geographically/linguistically diverse populations (9 Austroasiatic, 12 Austronesian, 8 Hmong-Mien, one Japonic, and one Koreanic, 12 Sinitic, 14 Tai-Kadai, 18 Mongolic, and 24 Tibeto-Burman), as well as 424 ancient genomes from 70 archaeologic sites or genetic groups from southern Siberia, Mongolia, China, Japan, Nepal and China. We first evaluated the genetic relationships among our included 1,498 ancient and present-day individuals using PCA. We identified two parallel genetic clines that were stretched out along PC2 (**Figure 1A**) in the two-dimensional plots: Northern East Asian genetic cline consisted of Mongolian ancients and modern Tungusic/Mongolic speakers at one end and Tibeto-Burman-speaking populations at the other; Southern East Asian genetic cline comprised Austronesian/Austroasiatic people at one end and inland Hmong-Mien speakers at the other. The third genetic cline located between southern Tai-Kadai and northern lowland Tibeto-Burman or Sinitic along PC1 linked the aforementioned northern and southern East Asian genetic clines, which is referred to as the Han Chinese related cline. This intermediate cline included our newly-genotyped Tanka people who were localized close with modern southern Han Chinese but showed a deviation toward inland Hmong-Mien-speaking Hmong and PaThen. A clear genetic landscape of substructure could be arranged along PC3 (**Figure 1B**): High-altitude Tibeto-Burman-speaking Tibetan and Sherpa were separated from others, and Austroasiatic people from the Mainland of Southeast Asia were separated from the island Austronesian. Here, Fujian Tanka people showed a genetic affinity with modern Austronesian and ancient Iron Age Hanben people. After excluded ancient populations from the Mongolian Plateau, the aforementioned genetic clusters/clines were visualized clearly via the top three components (**Figure 3C∼D**). Finally, we focused on populations from southern East Asia and Southeast Asia (**Figure 3E∼F**). Four genetic sub-clines (Hmong-Mien cline, mainland Austroasiatic cline, inland Austronesian cline, and northern mainland Sinitic cline) and one Tai-Kadai genetic cluster were identified. Tanka people were grouped closely with Sinitic speakers, not with southern Chinese indigenous populations. We also found that Xinshizhou Tanka has deviated from the Shacheng Tanka and southern Han Chinese.

**Figure 1.**
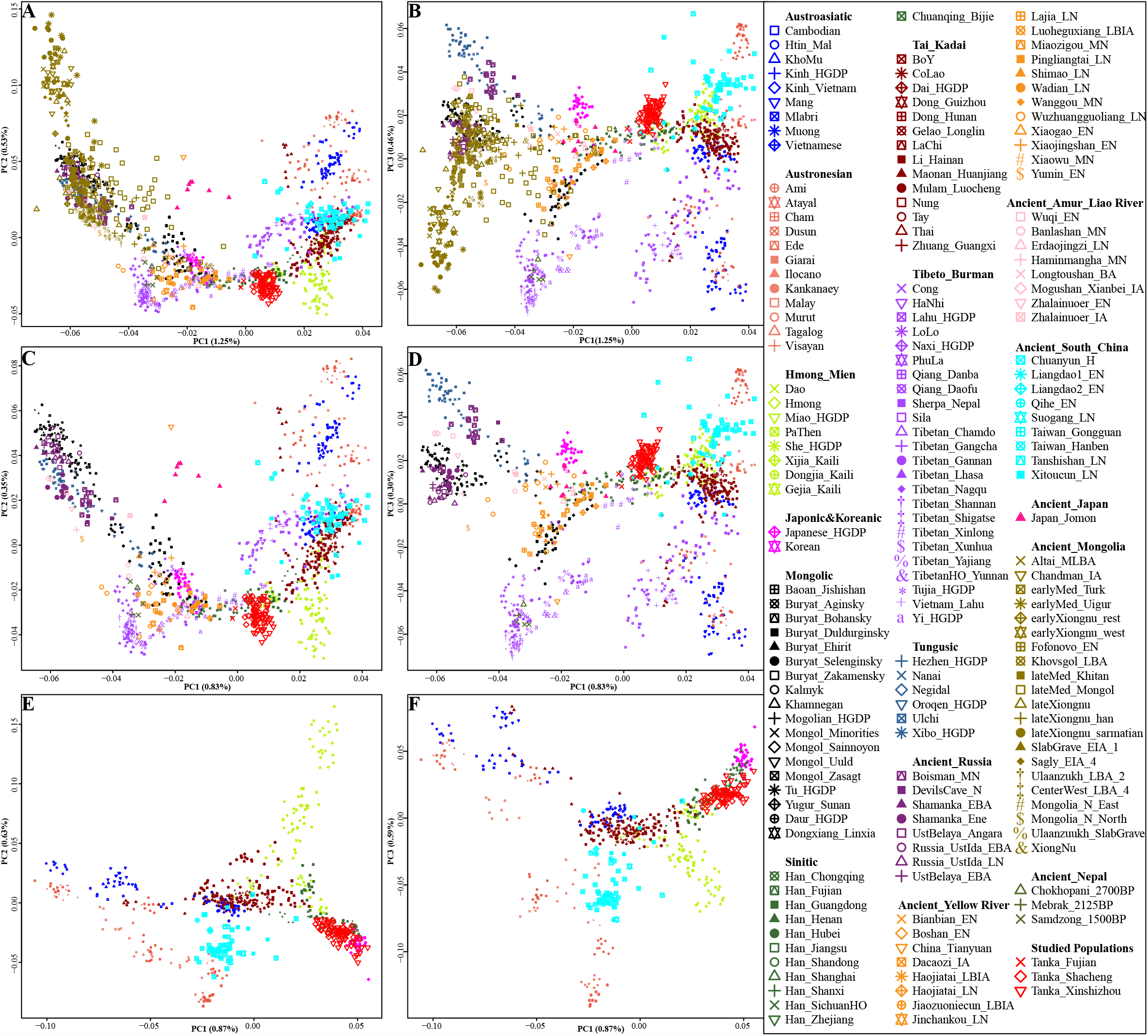
The general pattern of genetic relationship among modern and ancient eastern Eurasian via principal component analysis (PCA). (**A∼B**) Clustering results of all included modern and ancient eastern Eurasian populations based on the top three components. (**C∼D**). PCA results among eastern Eurasians after removing ancient Mongolian populations with western Eurasian admixture signals. (**E∼F**). Patterns of genetic affinity inferred from PCA focused on southern Chinese and Southeast Asian populations. Our included ancient reference populations were projected into the modern genetic framework. Modern populations were color-coded based on linguistic affinity and ancient populations were color-coded via geographical division.

Model-based clustering via ADMIXTURE among 1,498 Eurasian individuals from 178 modern and ancient individuals was conducted to further explore the genetic diversity and ancestry composition (**Figure S1∼2**). Cross-validation results showed that the model with eight predefined ancestral sources was the best-fitted one (**Figure 2**). Four northern East Asian ancestry components were identified respectively existed in Boisman_MN, Mongolia_N_North, Late-Xiongnu-Sarmatian and Mebrak_2125BP with a maximized proportion. Four southern East Asian ancestry components were respectively maximized in Taiwan_Gongguan, Hmong, Mlabri and Htin. Shacheng Tanka was modeled as the admixture result of 0.256 coastal Gongguan-related or Austronesian-related ancestry, 0.299 inland Hmong-Mien-related, and 0.320 high-altitude Mebrak-related ancestries. The other two studied populations also could be fit well with these three sources of ancestry: Hmong-Mien-related, Austronesian-related, and Tibeto-Burman-related, in respective proportions of 0.320, 0.274, and 0.311 (Xinshizhou Tanka); and 0.291, 0.215, and 0.338 (Fujian Han). Interestingly, we found a homogeneous Tanka-specific blue ancestry when the assumed ancestry sources increased (K≥9). Tanka-dominant ancestry played an important role in the Austroasiatic, Tai-Kadai, Hmong-Mien, and Sinitic speakers, which were rare or no proportion in Austronesian-speaking populations. Compared with Tanka people, geographically close Han harbored more Tibetan-dominant ancestry from North China.

**Figure 2.**
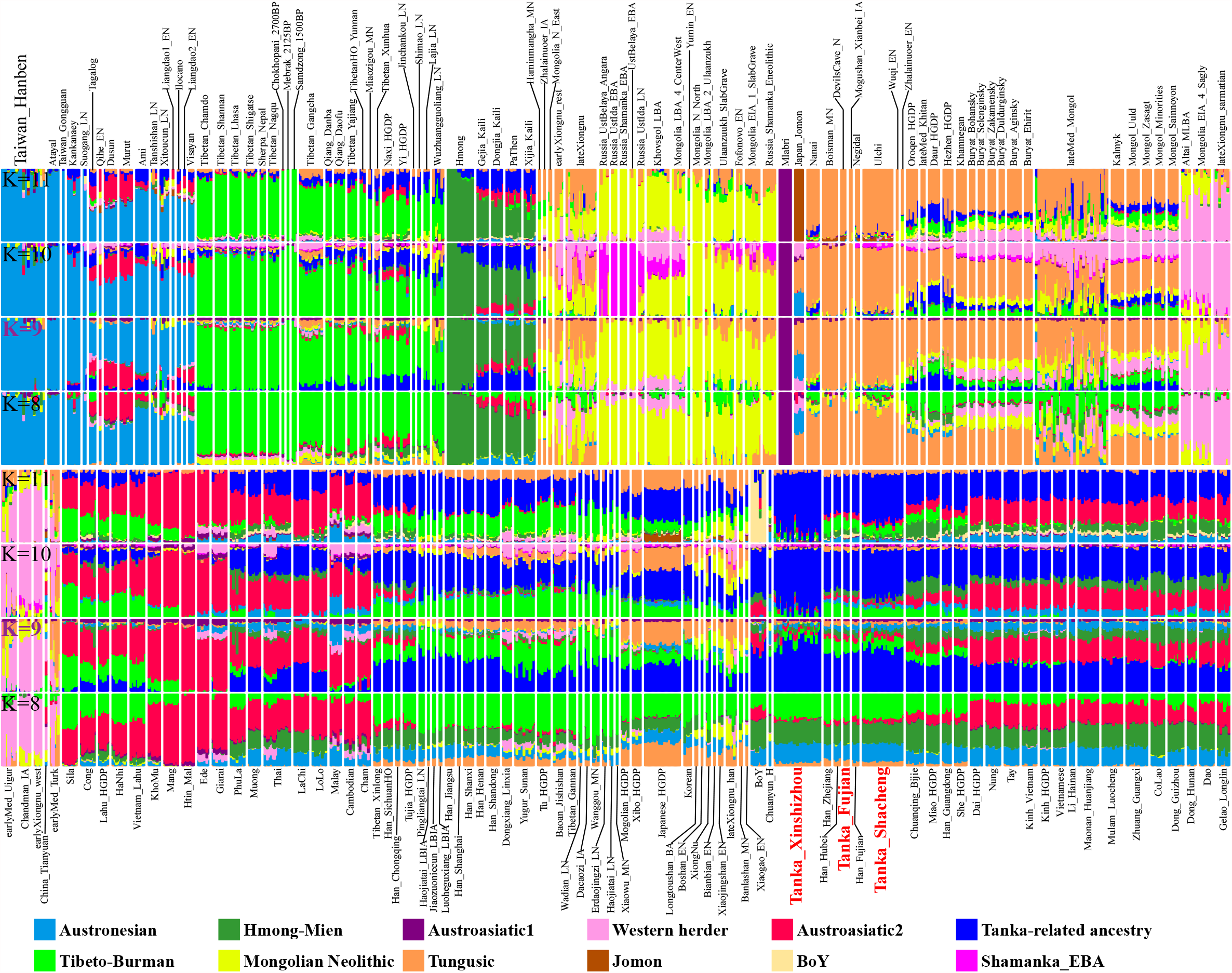
Model-based ADMIXTURE clustering analysis among eastern modern and ancient Eurasian populations. Predefined ancestral numbers of eight to eleven were chosen there. The full landscape of genetic structure was submitted in Supplementary Figure S3.

### The genomic affinity of Tanka people in the context of modern and ancient eastern Eurasian

We next calculated two statistical parameters for evaluating the genetic similarities and differences between Fujian Han and Tanka people relative to other modern and ancient eastern Eurasian reference populations. The unbiased pairwise genetic distance of Fst was measured between three populations and other 65 southern East Asians or Southeast Asians, the smaller value between Tanka people and Han Chinese populations indicated a closer genetic relationship (**Table S1**). We observed northern Shacheng Tanka possessed the smallest genetic distance with new-studied Fujian Han (0.0055), followed by Zhejiang Han (0.0059) and previously published Fujian Han (0.0061). For ancient southern coastal ancient populations, Shacheng Tanka harbored a closer genetic relationship with the Taiwan Iron Age Hanben population (0.0444), followed by the Late Neolithic Xitoucun (0.1776) and Tanshishan (0.1993). Southern Xinshizhou Tanka harbored the smallest genetic relationship with two Fujian Han Chinese populations among modern reference populations as well as possessed stronger affinity with Hanben and Late Neolithic Fujian populations among ancient reference groups. Newly-studied Fujian Han showed the most genomic affinity with all northern and southern Han Chinese than two geographically close Tanka groups, consistent with the clustering pattern of the previously published Fujian Han. Our result indicated that although a genetic affinity between Tanka people and Han Chinese could be identified, Tanka people have more shared genetic ancestry with southern indigenous Austronesian, Tai-Kadai and Hmong-Mien populations compared with southern Han Chinese. Shared genetic drift between three Fujian populations and 427 Eurasian modern and ancient populations was further evaluated via the outgroup-*f*_*3*_(Eurasian reference populations, three studied populations; Mbuti). Lager *f*_*3*_-values indicated a closer genetic affinity (**Figure 3A∼B**). After excluding the overlapping SNP loci less than 10000, we observed consistent patterns of genetic affinity with East Asians, in which Tanka people shared more genetic drift with southern Han Chinese (**Table S2**). Among ancient populations, Tankan people shared the strongest genetic drift not only with southern Iron Age Taiwan indigenous people (Hanben and Gongguan) and Vanuatu Neolithic populations but also with the northern YRB Neolithic-to-modern people (**Figure 2A∼B**), which suggested the strongest genetic link between North China, Southeast China and Vanuatu via the ancient genetic legacy of southeastern farmer elites

**Figure 3.**
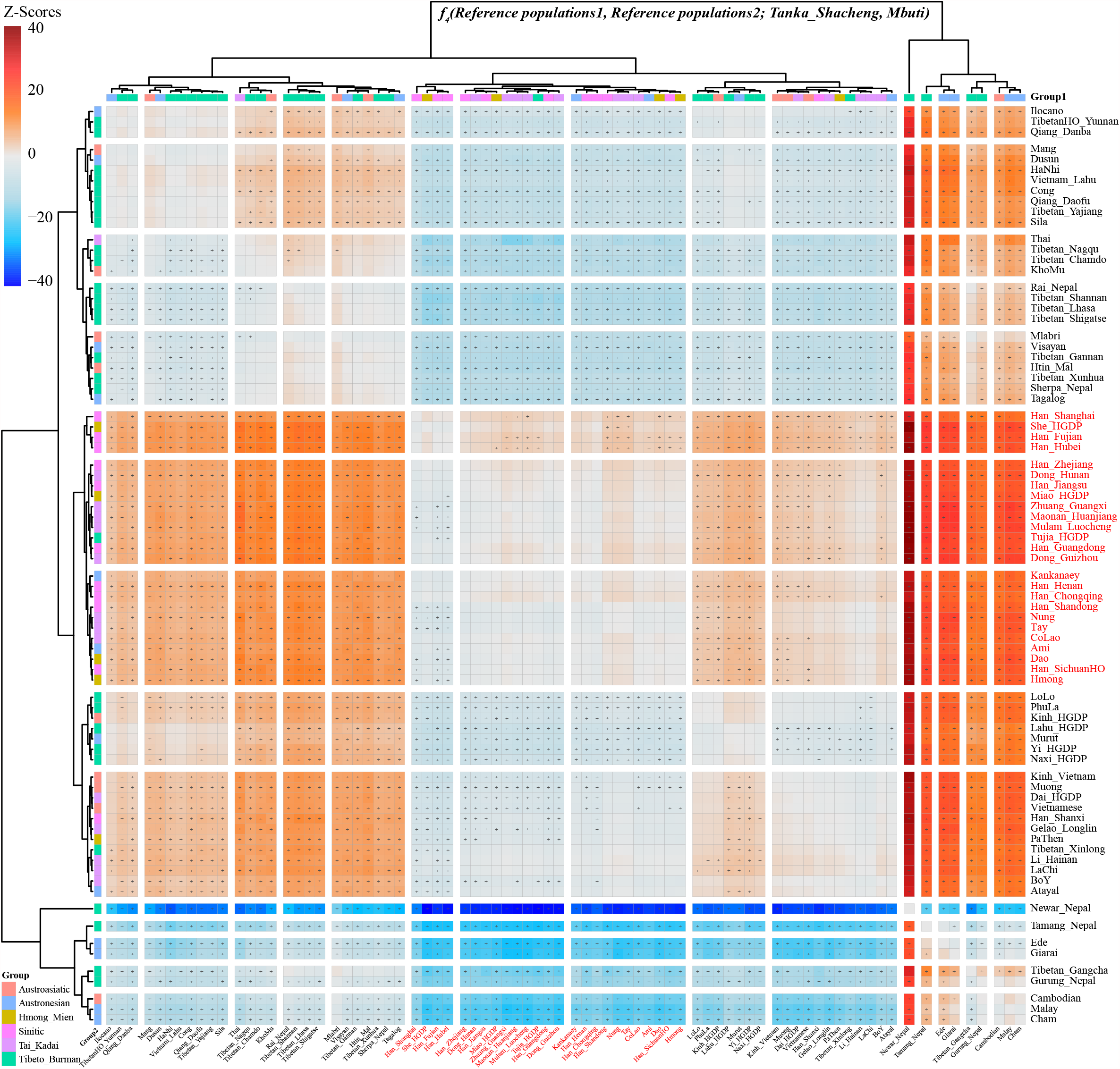
Genomic affinity and admixture signatures of our targeted populations. (**A**) Heatmap displayed the shared genetic between Shacheng Tanka and other eastern modern and ancient Eurasian populations. Red color denoted stronger affinity with Tanka people and green color denoted relatively far genetic relationship with Tanka speakers (**B**) Top twenty populations shared strong genetic affinity with Xinshizhou Tanka. The short bar showed the three-fold of standard error. (**C**) Admixture-*f*_*3*_-statistics in the form of *f*_*3*_*(Source1, Source2; Shacheng/Xinshizhou Tanka/Fujian Han)* showed the top twenty pairs for potential admixture sources. Red asterisk showed obvious statistically significant results with absolute Z-Scores larger than three. And green circle showed no statistically significant results.

Evidence for different genetic features of Tanka people relative to geographically close Han Chinese was further provided via admixture-*f*_*3*_(Source1, Source2; Focused populations). This type of allele-sharing-based *f*-statistics was widely used to explore the possible ancestral source candidates for one targeted population, in which statistically significant *f*_*3*_*-*values (Z-scores less than negative three) indicated obvious admixture signature. As shown in **Figure 3C**, we observed admixture signals in the new studied Fujian Han Chinese population with one source from northern East Asia associated with Tibeto-Burman-speaking populations (Tibetan and Qiang) and the other source candidates from southern East Asia related to Austronesian- or Tai-Kadai-speaking populations, such as *f*_*3*_*(Ami, Tibetan_Chamdo; Han_Fujian)= -5*.*26*SE*, which is consistent with the previous population genetic analyses focused on southern and central Han Chinese (G. He et al., 2020; G. L. He et al., 2020). We also found ancient northern East Asians, including the Early Iron Age SlabGrave people from Eastern Eurasian steppe, Neolithic Shimao, Jinchankou, Lajia, Wanggou and others combined with southern sources can produce negative *f*_*3*_-values, suggested the spatiotemporally northern East Asians contributed the genetic material into modern Fujian Hans. Similarly, southern ancient East Asians associated with Hanben, Tanshishan and Xitoucun combined with northern sources also could produce admixture signals for Fujian Han. However, we did not identify statistically significant admixture signatures in the admixture *f*_*3*_-statistics focused on Xinshizhou and Shacheng Tanka people, which is consistent with the observed homogenous genetic structure in our model-based ADMIXTURE result with ancestral sources larger than eight (**Figure 2**). We also observed 48 source pairs that possessed negative-*f*_*3*_-values in Shacheng Tanka and 10 pairs showed as negative-*f*_*3*_-values in Xinshizhou Tanka. These weak signals also provided clues that suggested more genetic interaction between Shacheng Tanka people with their geographical neighbors than it in Xinshizhou Tanka.

### Genetic continuity and admixture of southeastern Chinese Tanka people

We following validated the genetic homogeneity observed in PCA and ADMIXTURE results using *f*_*4*_-statistics. No statistical deviations were observed among *f*_*4*_*(Tanka_Shacheng, Tanka_Xinshizhou; Eastern Eurasian ancients, Mbuti)*, in which 75 ancient populations were used here, suggesting the genetic cladality between two geographic different Tanka populations. If we settled the threshold of the absolute Z-score as two, we could find the status that northern East Asians shared more alleles with geographically northern Shacheng Tanka relative to the southern Xinshizhou Tanka people, such as Neolithic Miaozigou people with *f*_*4*_*(Tanka_Shacheng, Tanka_Xinshizhou; Miaozigou_MN, Mbuti)=2*.*401*SE* and Iron Age Zhalainuoer people with *f*_*4*_*(Tanka_Shacheng, Tanka_Xinshizhou; Zhalainuoer_IA, Mbuti)=2*.*90*SE*. Similarly, we observed Xinshizhou Tanka harbored more Hanben-related ancestry compared with Fujian Han. Compared with 134 modern populations, Austronesian speakers shared more derived alleles with two Tanka people relative to Fujian Han with the negative Z-scores smaller than -2 in *f*_*4*_*(Han_Fujian, Tanka_Xinshizhou/Tanka_Shacheng; Austronesian speakers, Mbuti)*. Summarily, Tanka people shared more southern East Asian ancestry related to Austronesian or Tai-Kadai-speaking populations compared with southern Han Chinese, suggesting the different genetic history between Tanka and their neighbor of Hans.

We further explored the genomic affinity between three newly-genotyped populations between other ancient and present-day eastern Eurasian populations via *f*_*4*_*(208Eurasian1, 208Eurasian2; three studied populations, Mbuti)*. Compared with northern East Asians, three Fujian populations shared a more common genetic ancestry component related to modern southern East Asian, Southeast Asian and modern Sinitic speakers, as statistically negative *f*_*4*_-values were observed in *f*_*4*_*(Northern East Asians, Southern East Asians; Tanka or Han in Fujian, Mbuti)*. Focused on southern populations (**Figure 4 and S4**), strong genetic affinity was observed between northern Shacheng Tanka and four populations (Han_Shanghai, She, Han_Fujian and Han_Hubei), followed by other Sinitic speakers, Tai-Kadai, Hmong-Mien-speaking Miao, Austronesian-speaking Ami and Kankanaey and lowland Tibeto-Burman-speaking Tujia. Xinshizhou Tanka showed a similar pattern of genomic affiliation (**Figure S4**). Compared with inland southern East Asians, Tanka people shared more ancestry with southeastern coastal East Asians, especially for Austronesian speakers with statistically negative values in *f*_*4*_*(Dai, Ami; Tanka_Shacheng, Mbuti)*. Compared with Bronze Age steppe pastoralists (Afanasievo and Sintashta), Tanka people shared most derived alleles with southern Hanben and northern late medieval Mongolian people. Compared with Fujian Late Neolithic Tanshishan population, three studied populations shared more alleles with the Late Neolithic Longshan people from the Yellow River basin due to the observed positive values in *f*_*4*_*(Yellow River farmers, Tanshishan; Tanka, Mbuti)* (Z-Scores: Tanka_Xinshizhou, 2.432; Tanka_Shacheng, 2.447; Tanka_Fujian, 2.627). Results from the affinity-*f*_*4*_ statistics showed Tanka people shared more alleles with Han Chinese populations compared with southern Chinese minority populations. Our findings also showed that Tanka people shared more alleles related to the southern Chinese Tai-Kadai or Austronesian people compared with southern Han Chinese, suggesting that the Tanka people not only had a strong genetic affinity with Han people but also harbored additional gene flow from surrounding indigenous populations.

**Figure 4.**
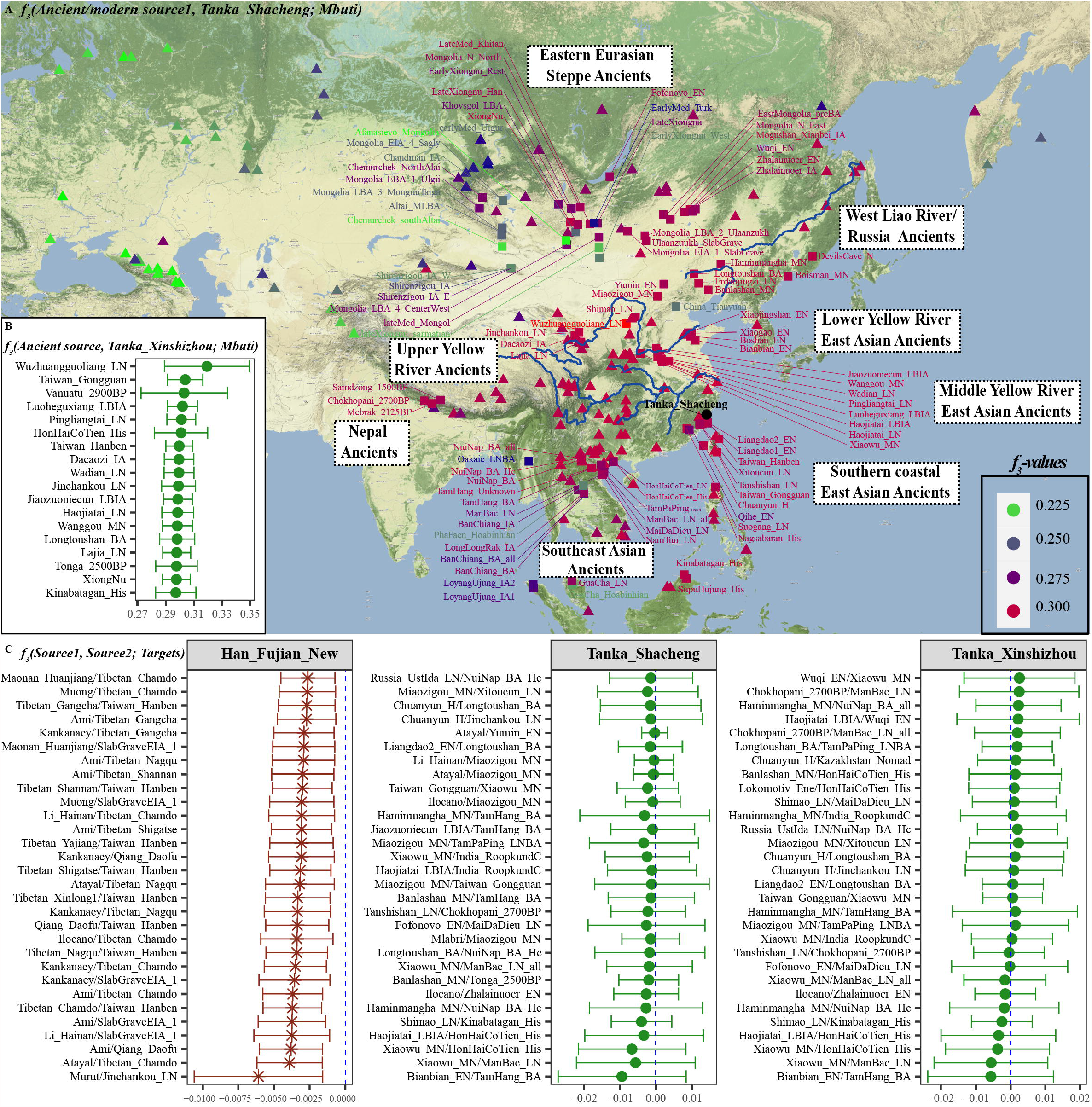
A formal test of genomic affinity in Shacheng Tanka people inferred from the two-population comparison *f*_*4*_-statistics in the form *f*_*4*_*(Reference population1, Reference population2; Tanka_Shacheng, Mbuti)*. Red color denoted the positive *f*_*4*_-values, which suggested Shacheng Tanka people shared more derived mutations with reference population1 (left population lists), and blue color showed the negative *f*_*4*_-values, which suggested Shacheng Tanka people shared more alleles with reference population2 (bottom population lists). Statistically significant results were marked with the ‘+’.

Additionally, to explore the genetic contribution to modern Tanka’s gene pool, we performed *f*_*4*_*(Reference populations, Tanka people; Southern modern and ancient populations, Mbuti)*. If we included southern modern and ancient populations as the direct ancestors or their ancestral proxies, significant negative *f*_*4*_-values would be expected observed. As showed in **Figure 5 and S5**, signatures of genetic contributions (blue color indicated genomic affinity with Tanka people) were observed not only when we assumed southern East Asians as the Tanka’s ancestral sources but also identified when we assumed northern modern Tibeto-Burman speakers and northern East Asians as their ancestral contributors. When we hypothesized northern Neolithic-to-Iron Age East Asian people as their ancestor (Northern East Asian origin hypothesis), we also identified too many blue signals evidenced for the genetic contribution from southern East Asians. These identified shared alleles suggested a strong phylogenetic correlation between Tanka people and both northern and southern East Asians. To further validated some unique ancestral sources directly contributed to the Tanka people, we analyzed *f*_*4*_-statistics in the new form *f*_*4*_*(Ancestral sources, Tanka people; reference populations, Mbuti)*. Non-*f*_*4*_-values showed the statistical significance (Absolute Z-Scores larger than three) were identified in *f*_*4*_*(Chuanyun_H/Miaozigou_MN/Haojiatai_LN/Tujia_HGDP/Han_Chongqing/Han_Fujian/Han_Zhejia ng, Tanka people; Reference populations, Mbuti)* showed the genomic affinity between Tanka people and southern Han Chinese as well as the Yellow River late Neolithic ancient of Haojiatai people (forming one genetically indistinguishable clade), which was consistent with the expected patterns if their northern China Origin is the true history model. Following, we used other YRB millet farmers as their director ancestors, we identified additional admixture signatures from southern East Asians in *f*_*4*_*(Jiaozuoniecun_LBIA/Jinchankou_LN/Wadian_LN/Xiaowu_MN/Haojiatai_LBIA/Pingliangtai_LN, Tanka people; southern Chinese populations, Mbuti)*, suggesting additional mixture events occurred in the past, which is consistent with the identified marginal negative-*f*_*3*_-values in admixture-*f*_*3*_-statistics. Similar patterns were identified when we used the southern East Asians as their direct ancestor, suggesting additional gene flow contributed to the formation of the Tanka people also from northern East Asians. Formal tests in *f*_*4*_-statistics demonstrated both northern East Asians and southern East Asians participated in the formation of the Tanka people, and it also possessed more southern indigenous ancestry related to Austronesian or Tai-Kadai speakers compared with Fujian Han.

**Figure 5.**
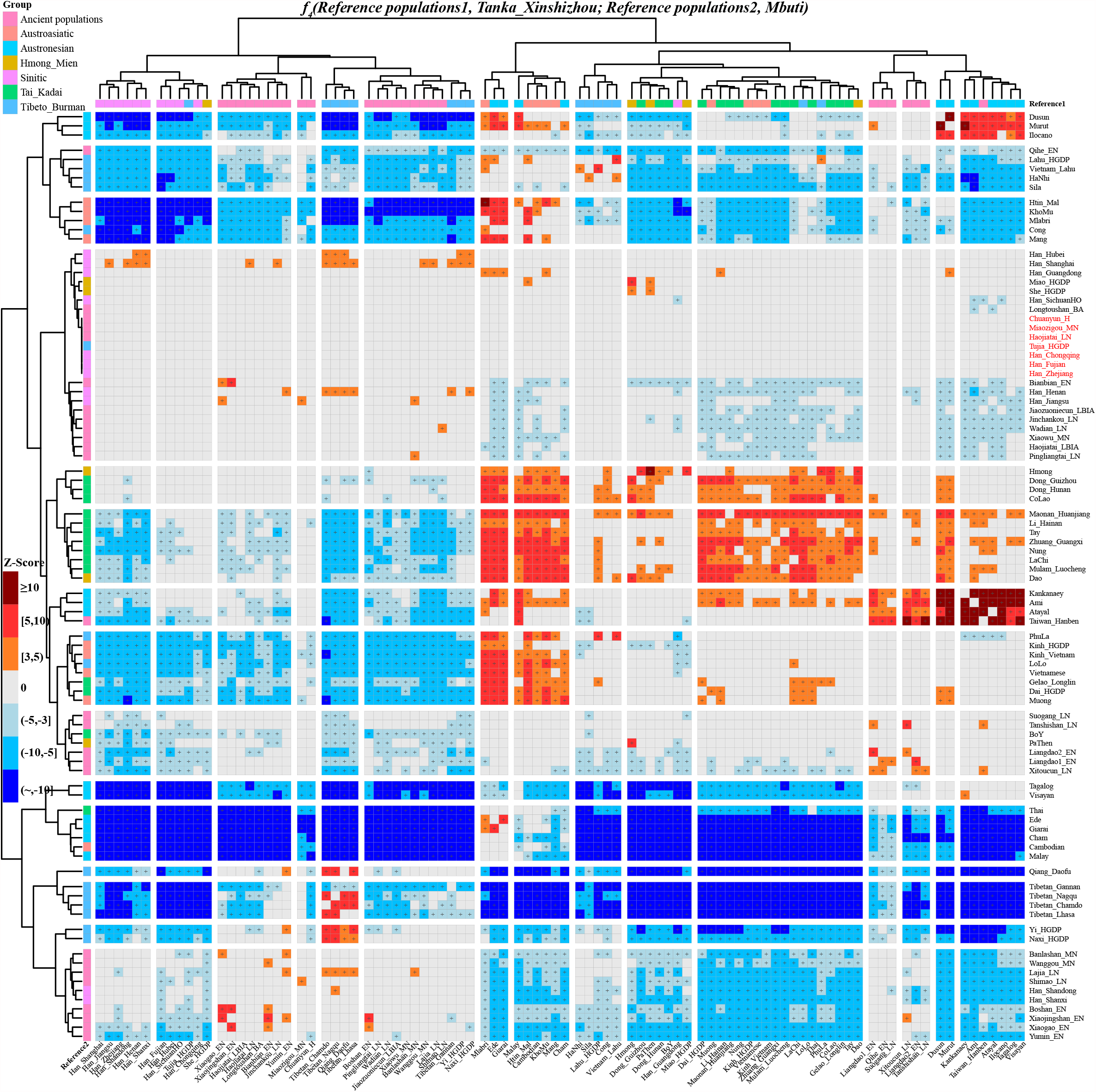
A formal test of genomic continuity and admixture in Xinshizhou Tanka people inferred from the two-population comparison *f*_*4*_-statistics in the form *f*_*4*_*(Reference population1, Reference population2; Tanka_Xinshizhou, Mbuti)*. Red color denoted the positive *f*_*4*_-values, which suggested Reference population2 (bottom population lists) shared more derived mutations with reference population1 (left population lists), and blue color showed the negative *f*_*4*_-values, which suggested reference population2 shared more alleles with Xinshizhou population and gray color showed no statistically significant results were observed. Statistically significant results were marked with the ‘+’.

### Phylogenetic history reconstruction and quantification of the mixed ancestry

We constructed one unrooted phylogenetic framework (**Figure 6A**) among southern Chinese Hmong-Mien, Tai-Kadai, Austroasiatic, and Austronesian modern people, as well as ancient southern coastal East Asians (Neolithic Tanshishan and Xitoucun, and Iron Age Hanben). PCA results among TreeMix-used populations showed three genetic clines among these included populations: Austronesian and ancient Taiwanese-related, Austroasiatic-related and others (**Figure 6B**), consistent with the patterns in previous southern PCA with ancient samples projected. Tanka people were clustered closely with Han Chinese but not with Tai-Kadai or Austronesian speakers. In accordance with the genetic clustering in ADMIXTURE and PCA results, four branches were identified in the TreeMix-based phylogenetic framework. Hmong-Mien branch was clustered close with the Sinitic branch, which diverged from the common ancestor of Tai-Kadai, Austronesian and Austroasiatic speakers. Three studied populations were grouped closely and firstly clustered with neighboring Han (Fujian and Guangdong) and Chuanqing and She people. To further validated the Sinitic-affinity or North China Origin hypothesis of primary ancestry of Tanka people, we reconstructed the phylogenetic trees based on the *f*-statistics (*f*_*2*_, *f*_*3*_ and *f*_*4*_) using qpGraph. As shown in **Figure 7**, we used Neolithic populations from Mongolia Plateau (MNE), Qinghai-Tibetan Plateau (Chokhopani) and Yellow River Basin (Xiaowu_MN) as the ancient northern East Asian source, and used mainland late Neolithic Tanshishan and Iron Age Hanben as the ancient southern East Asian source to construct the basic phylogenetic framework. Tanka people could be modeled as the main ancestry from ancient northern East Asians: Xinshizhou Tanka derived 91% of their ancestry from Xiaowu Yangshao millet farmer and the reminding from the deep diverged eastern Eurasian related to Onge in the deep admixture model (**Figure 7A**), and it can be also modeled via 64% Xiaowu-related ancestry and 36% Tanshishan-related ancestry in the recent admixture model (**Figure 7B**). We further used different northern and southern sources to estimate the date of these admixture events via the decay of the linkage disequilibrium (**Figure 7C∼D**). We found that major admixture events mainly occurred during the Late Neolithic to medieval times with different predefined ancestral sources, which was consistent with the estimated dates from GLOBETROTTER.

**Figure 6.**
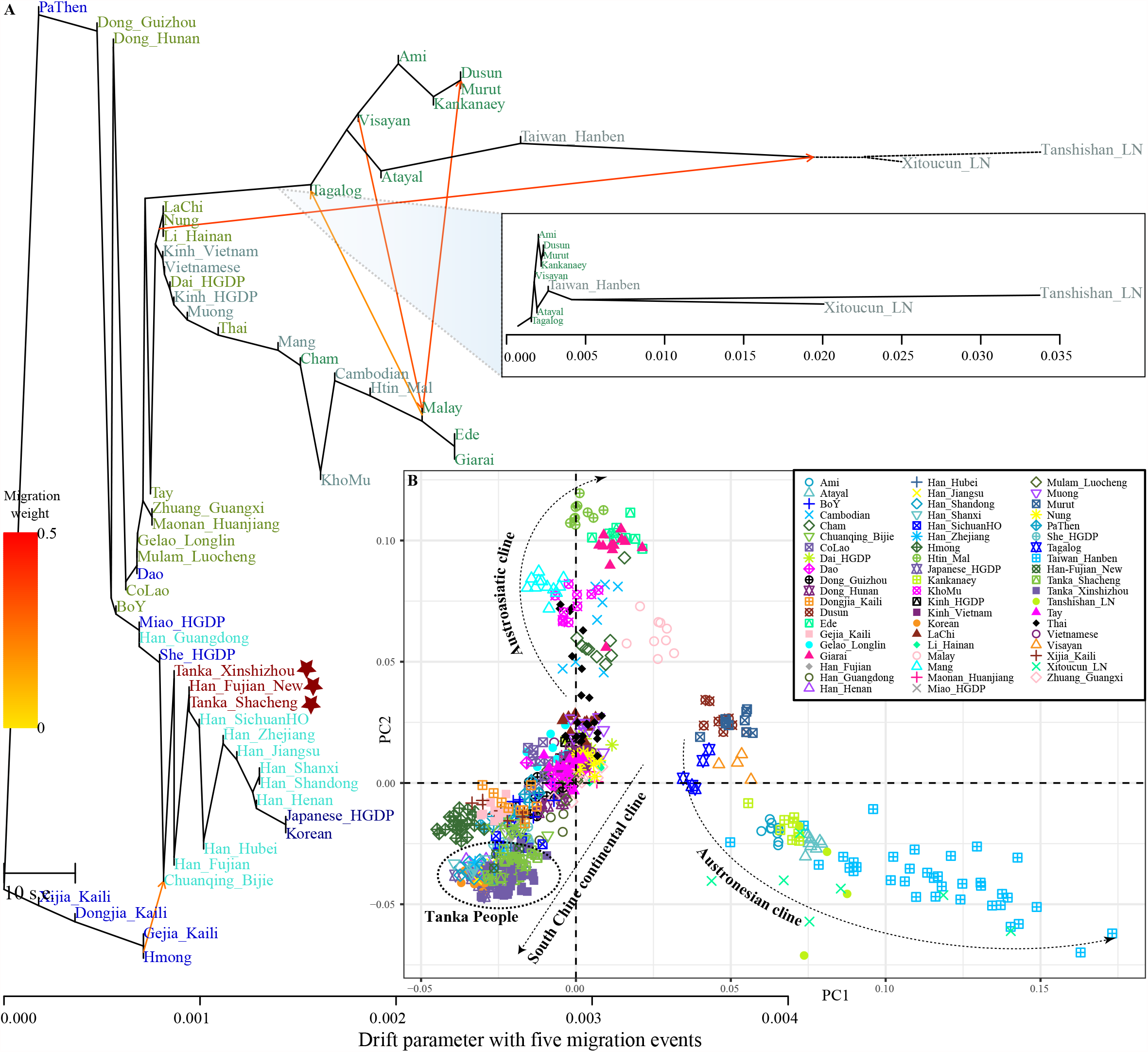
The TreeMix-based phylogenetic tree showed the genetic relationship between modern and ancient East Asians. (**A**) Tree with four assumed admixture events. (**B**) Population relationships among TreeMix-used populations without ancient populations were projected.

**Figure 7.**
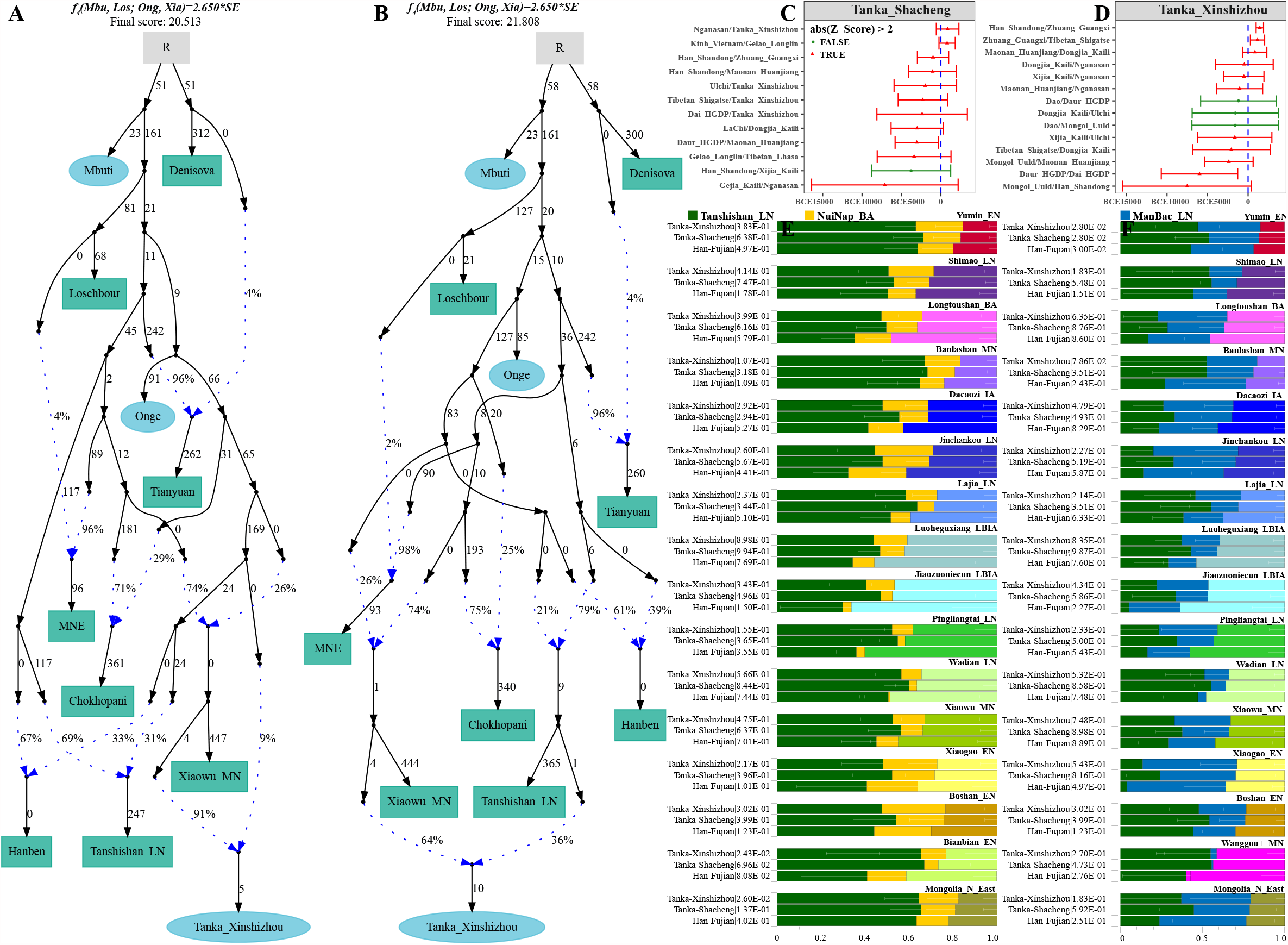
Genetic drift-based phylogenetic phylogeny showed population split and gene flow events. (**A**) The targeted population was modeled as derived their ancestry from the Yellow River Yangshao lineage and one ancient southern deep diverged eastern Eurasian lineage. (**B**). Tanka people were modeled as the admixture of two Neolithic East Asian lineages. Genetic drift was marked as 1000 times of *f*_*2*_ values. Dot blue lines denoted the admixture events and corresponding admixture proportions were also marked. (C∼D). Dates of admixture events estimated via ALDER. (**E∼F**) QpAdm results showed the admixture proportion of three-way admixture models. Admixture proportion was estimated used the northern East Asian-Tanshishan_LN-NuiNap_BA three-way admixture model (**E**). Admixture proportion was estimated when we used the Neolithic ManBac people as the inland southern East Asian(**F**).

Finally, we used *qpAdm/qpWave* and the three-way admixture model to evaluate the fine-scale ancestral proportion from the southern sources. Here, we used late Neolithic Tanshishan people as the proxy of southern coastal ancestral populations and used Bronze Age NuiNap and late Neolithic ManBac from Vietnam as the proxies of inland ancestral sources. Two main findings we identified from *qpAdm* results (**Figure 7E∼F**). Firstly, all included three modern populations could be successfully fitted via at least one of we provided mixed models, even including the Han Chinese in model of Tanshishan-NuiNap-Jinchankou, suggesting the complex ancestral sources of modern southern Han Chinese, which is differentiated from the demographic history of present-day northern Han people. Secondly, compared with southern Han, Tanka people harbored less northern East Asian-related ancestry, and Xinshizhou Tanka people possessed more inland southern East Asian-related ancestry and less northern East Asian-related ancestry.

### Demographic history of Tanka people inferred from the sharing haplotype patterns

We subsequently explored the genetic similarities and differences within and between Tanka people and their adjacent East Asians(Chen et al., 2021; He et al., 2021; Y. Liu et al., 2021; Lu et al., 2020; Jin Sun et al., 2020; Q. Wang et al., 2020; Yao et al., 2021) based on the genetic variants of denser SNP data (approximately 700K). we evaluated the pairwise Fst genetic distances again based on the denser SNP dataset and found a closer relationship between Shacheng Tanka and Fujian Tanka (0.0062) than it between Xinshizhou Tanka and Fujian Tanka people (0.0100), all these less than the genetic differences between Shacheng and Xinshizhou Tanka people (0.0130). We conducted PCA analysis among three Tanka populations and found population substructures among them. We could identify three branches (one Shacheng and two Xinshizhou groups): Fujian Tankas were clustered between Xinshizhou and Shacheng (**Figure 8A**). Although the population substructure was identified here, no statistically *f*_*4*_ values were identified via f4(Xinshizhou1, Xinshizhou2; Reference populations, Mbuti). Thus, one population label was used in the frequency-based analysis.

**Figure 8.**
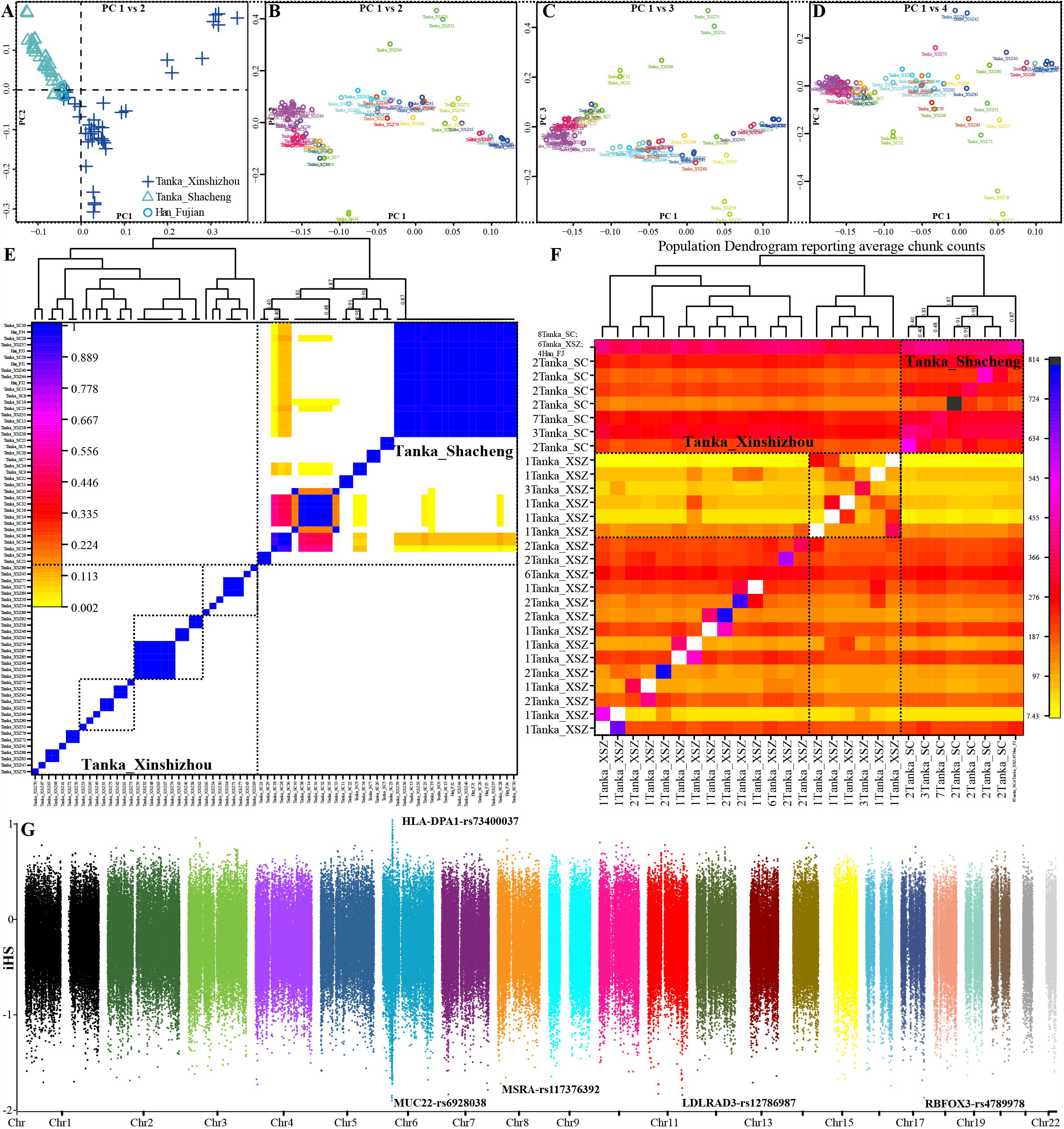
Finer-scale population structure within Fujian Tanka people using the phased haplotype data. (**A**). Plink-based PCA showed the substructure within Tanka people based on the allele frequency spectrum. (**B∼D**). Haplotype-based PCA showed the substructures within Tanka people. (E∼F) The pairwise coincidence and heatmap showed the individual clustering pattern. (G) Natural selection signatures inferred from the estimated iHS.

We following phased the successive independent SNP data to haplotype form from maternal and paternal sides based on the haplotype phased statistical methods. Fine-scale population structures were further comprehensively characterized based on the haplotype data. PCA results based on the coancestry matrix also confirmed the aforementioned population substructure but with subtle differences for substructures (**Figure 8B∼D**). We observed the genetic homogeneity within Shacheng people and genetic heterozygosity within Xinshizhou people via the scattered plots in the two-dimensional plots. We subsequently explore the patterns of shared haplotype based on the ChromoPainter and FineSTRUCTURE. We evaluated the demographic parameters based on the random extracted samples (including the effective population size of Tanka people (Ne: 290.776) and mutation rate (mu: 0.0006513)). The inferred population affinity and dendrogram based on the pairwise coincidence showed that eight Tankas from Shacheng, six Tankas from Xinshizhou and other 20 Shacheng Tanka people clustered together and formed one genetically homogeneous population. The remaining Xinshizhou Tankas people formed the other branch (**Figure 8E∼F**). We also calculated the iHS values based on phased haplotypes and identified some natural selection signatures from Chromosomes 6, 8, 11 and 17, which respectively associated with the genes of HLA, MUC22, MSRA, LDLRAD3 and RBFOC3

We finally explored the genetic relationship between Tanka people with other 29 Chinese populations based on the shared haplotypes. Plink-PCA based on the allele frequency spectrum among 32 populations revealed two genetic clines consisted of Guizhou Hui people and Yunnan populations, which were located distinct from others (**Figure 9A**). Tanks people clustered together and also showed a distinct genetic relationship with geographically close Guizhou populations but close with Shaanxi Hans, which further confirmed in the PCA patterns based on the coancestry matrix (**Figure 9B**). Population dendrogram based on the pairwise coincidence and the TreeMix-based phylogenetic relationship showed a close genetic relationship between Fujian Tanka and Fujian Han, as well as a close genetic relationship with northern Hans from Shaanxi than inland Guizhou and Yunnan minorities (**Figure 9C∼E**). Results from the pairwise IBD showed Fujian Han harbored the longest shared IBD with Tanka from Xinshizhou (20.373) and Shacheng (19.571), following by Ankang Hans, Chuanqing and other Shaanxi populations. Tanka people not only shared the longest IBD within populations but also shared relatively longer IBD with both northern Hans and southern Hmong-Mien speakers, suggesting Tanka people received gene fluence from both northern and southern East Asians (**Figure 9F**). Pairwise Fst also showed a close relationship between Fujian populations with Chongqing Miao and Tujia and northern Shaanxi Hans. Clustering patterns based on the Fst matrix showed Fujian Han clustered with other Han populations, but Tanks people clustered with Guizhou Manchu, Mongolian and other Hmong-Mien-speakers (**Figure 9G**).

**Figure 9.**
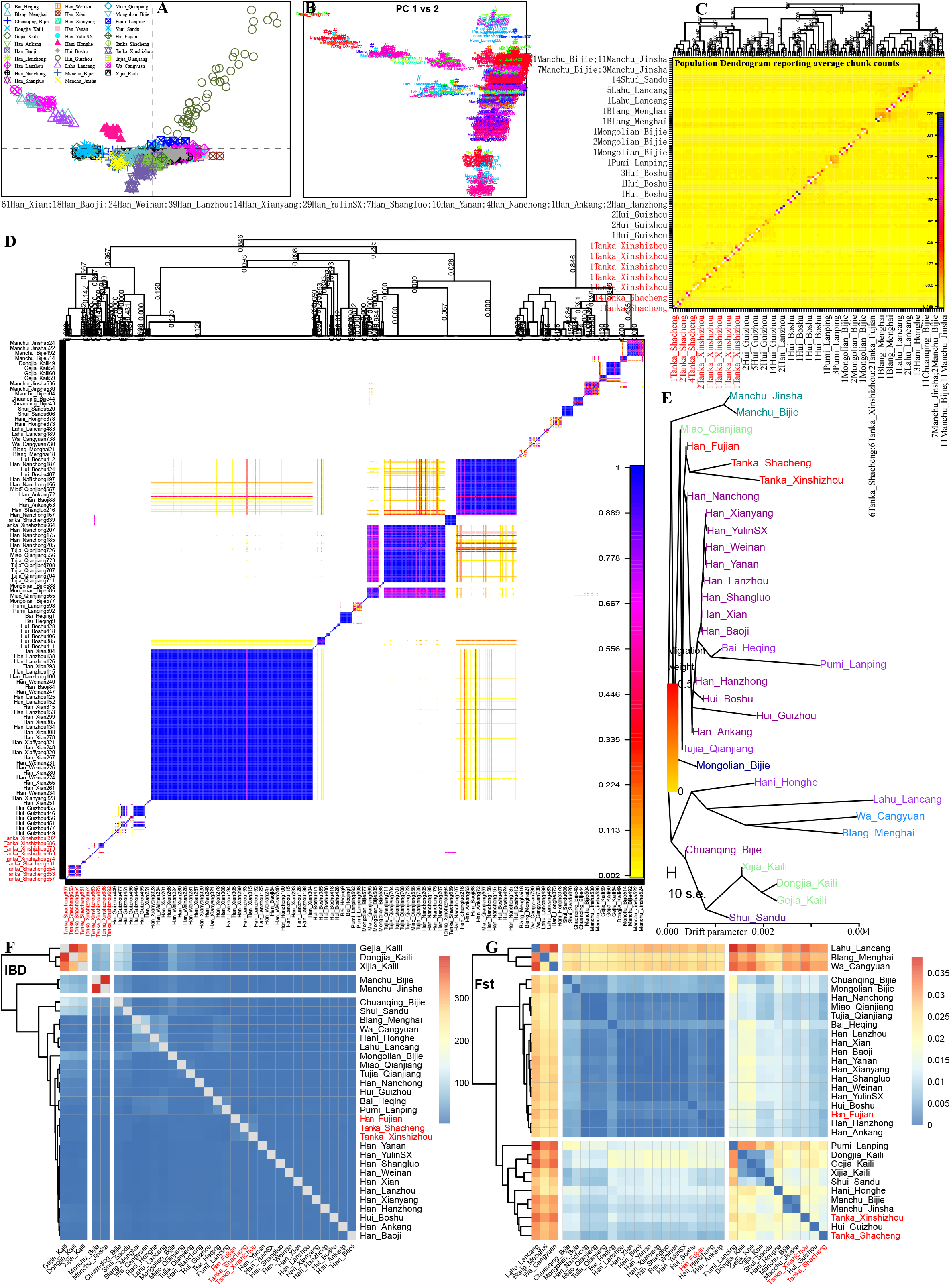
Finer-scale population structure between Fujian Tanka people and other 39 Chinese populations using the phased haplotypes. (**A∼B**). Patterns of the genetic relationship inferred from PCA based on the allele frequency spectrum and haplotype-based coancestry matrix. (**C∼D**) FineSTRUCTURE results showed population structure based on the shared number of the haplotype chunks. (E). The TreeMix-based phylogenetic tree showed the genetic relationship among 42 Chinese populations. (F∼G) Heatmap visualized based on the pairwise Fst and IBD matrixes.

### Uniparental genetic legacy

We genotyped paternally and maternally phylogenetic information SNPs in Tanka males at a higher resolution. Overall, the main founding paternal lineage in Tanka people in this study were O1a1a1a1a1a1-M119-F492, O1b1a1a1a1b1b-M95-CTS651, O1b1a2a1-Page59, O2a1c1a1a1a1d1-F325-F930, and O2a4b1a1a1.These observed paternal haplogroup types are also the main patrilineal types of people in southern China, but the patrilineal composition between the two groups is quite different. Lineages of O1a1a1a1a1a1-F492 and O1b1a1a1a1b1b-M95-CTS651 exited in Shacheng Tanka with a high proportion (50%, 17/34), but these two types do not exist in Xinshizhou Tanka. Similarly, O2a1c1a1a1a1d1-F325-F930 and O2a2b1a1a4a-F5-Z25853 accounted for a high proportion of Xinshizhou Tanka people (36%,14/39), but they did not exist among Shacheng Tanka people. Haplogroup O1b1a2a1-M268-P59 is the only shared patrilineal type among the two groups. Patriline O1a--M119 is one of the main patrilineal types of Austronesian and Tai-Kadai speakers, and there is also a certain proportion in geographically close Han populations. O1b1a1a1a1b1b-M95 is the major patrilineal type of the Hmong-Mien, Tai-Kadai, and Austroasiatic speakers, and it also has a high proportion in the Han ethnic group in South China. The downstream branch of O2-M122 is the main patrilineal type of the Han people. In short, the patrilineal genetic structure of the Tanka population observed here shows a strong association between the Tai-Kadai-speaking population and the southern Han populations. In terms of maternal lineage, the six main haplogroup types are identified, including M7b (21.9%, 16/73), M7c (9.59%, 7/73), F1a (9.59%, 7/73), FxF1a (8.22%, 6/73), D4 (8.22%, 6/73), M9a1a1 (6.85%, 5/73). Among these types, F1a and FxF1a have a higher proportion in the southern indigenous groups, while the other types have a higher proportion in the Han populations. In short, although the patrilineal lineages of the Tanka also show a significant connection with the Austronesian and Tai-Kadai groups in south China, it has a higher genetic similarity with the matrilineal lineages of the Han people.

## CONCLUSION

We generated genome-wide autosomal, paternal and maternal data of Tanka people to resolve the controversial hypotheses of the origins of Tanka People (Han origin, Ancient Baiyue origin, or admixture origin). We provided robust genetic evidence for the admixture original hypothesis of Tanka people based on the comprehensive population genetic reconstruction. Paternal genetic evidence found that Y-lineage haplogroup O1-M119 lineage reached a high frequency (35.2%,12/34) in Shacheng Tanka, which is higher than it in other southern Chinese Han populations. But this patrilineal genetic composition in Xinshizhou Tanka was almost the same as that of the southern Han. In the maternal genetic structure, a higher proportion of southern dominant maternal types could be observed. From the genetic variations from the autosomal genome-wide data, PCA and model-based ADMIXTURE results showed that Tanka people had their unique genetic structure, but kept a close relationship with geographically close southern Han Chinese populations. Shared genetic drift revealed from the outgroup-*f*_*3*_-statistic and admixture-*f*_*3*_-statistics further not only showed a stronger Han Chinese affinity but also displayed the marginal admixture signatures from the sources from southern and northern China, in which Tanka people could be modeled as the major ancestry related to the northern East Asian and minor ancestry related to Tai-Kadai-related populations. Thus, our results from the genome-wide data supported that Tanka people gave rise from the admixture between southward migration Han Chinese and southern indigenous people.

## Acknowledgments

We are grateful to all sample donors. This study was supported by the National Natural Science Foundation of China (31222030), China Postdoctoral Science Foundation (2021M691879). The funders had no role in study design, data collection and analysis, decision to publish, or preparation of the manuscript.

## Data availability

The raw genome-wide data can be obtained via corresponding authors.

## Legends of Figures

Figure S1. Geographical positions of two Tanka populations and one Han Chinese population collected from Fujian province in southeastern China.

Figure S2. Cross-validation error in the model-based ADMIXTURE analyses. The best model is the eight-source-based mixed model with the smallest cross-validation error (0.5750).

**Figure S3. Results of model-based ADMIXTURE analyses results with the predefined ancestral sources ranging from two to twenty**.

**Figure S4. Formal test of genomic affinity in Xinshizhou Tanka people inferred from the two-population comparison *f***_***4***_**-statistics in the form *f***_***4***_***(Reference population1, Reference population2; Tanka_Xinshizhou, Mbuti)***. Red color denoted the positive *f*_*4*_-values, which suggested Xinshizhou Tanka people shared more derived mutations with reference population1 (left population lists), and blue color showed the negative *f*_*4*_-values, which suggested Xinshizhou Tanka people shared more alleles with reference population2 (bottom population lists). Statistically significant results were marked with the ‘+’.

**Figure S5. A formal test of genomic continuity and admixture in Shacheng Tanka people inferred from the two-population comparison *f***_***4***_**-statistics in the form *f***_***4***_***(Reference population1, Reference population2; Tanka_Shacheng, Mbuti)***. Red color denoted the positive *f*_*4*_-values, which suggested reference population2 (bottom population lists) shared more derived mutations with reference population1 (left population lists), and blue color showed the negative *f*_*4*_-values, which suggested reference population2 shared more alleles with Shacheng population and gray color showed no statistically significant results were observed. Statistically significant results were marked with the ‘+’.

**Figure S6. Genetic drift-based phylogenetic phylogeny showed population split and gene flow events for Shacheng Tanka**. Tanka people were modeled as the admixture of two Neolithic East Asian lineages. Genetic drift was marked as 1000 times of *f*_*2*_ values. Dot blue lines denoted the admixture events and corresponding admixture proportions were also marked.

## Legends of Tables

Table S1. Pairwise Fst genetic distance between three studied populations and other 65 reference populations.

Table S2. Shared genetic drift between studied Tanka people and Han and other reference populations.

Table S3. Admixture signatures of admixture-f3 statistics focused on the new-studied Fujian Han population.

Table S3. Genetic homogeneity examined via the symmetric-f4(Studied population1, Studied population2; reference population, Mbuti).

