## Supplementary Figures for "The genomic formation of Tanka people, an isolated “Gypsies in water” in the coastal region of Southeast China"

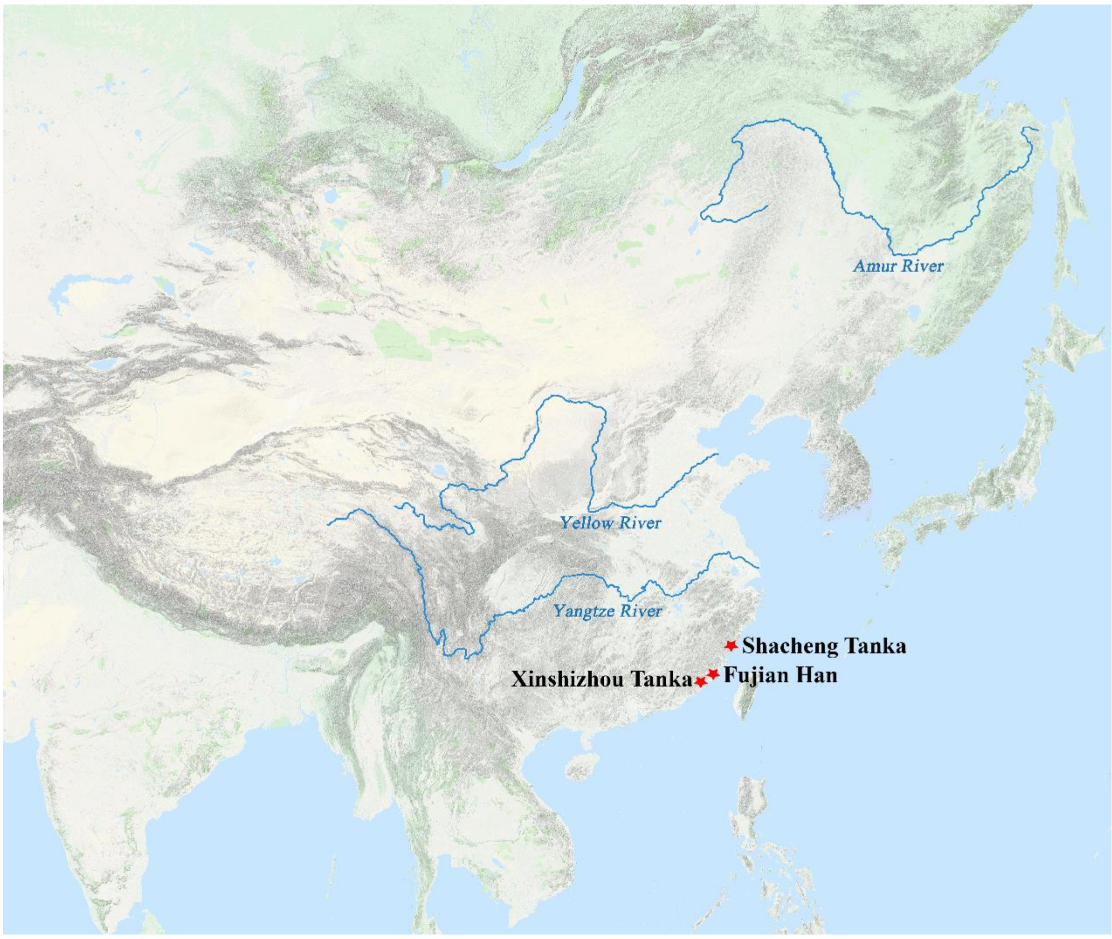


Figure 1. Geographical positions of two Tanka populations and one Han Chinese population collected from Fujian province in southeastern China.

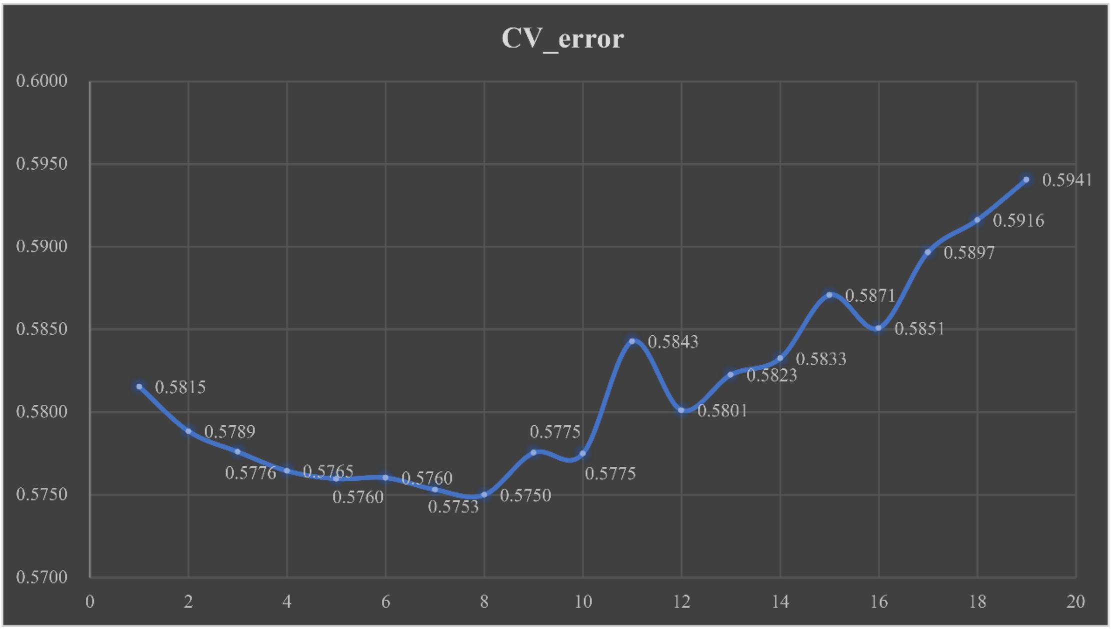


Figure S2. Cross-validation error in the model-based ADMIXTURE analyses. The best model is the eight-source-based mixed model with the smallest cross-validation error (0.5750).

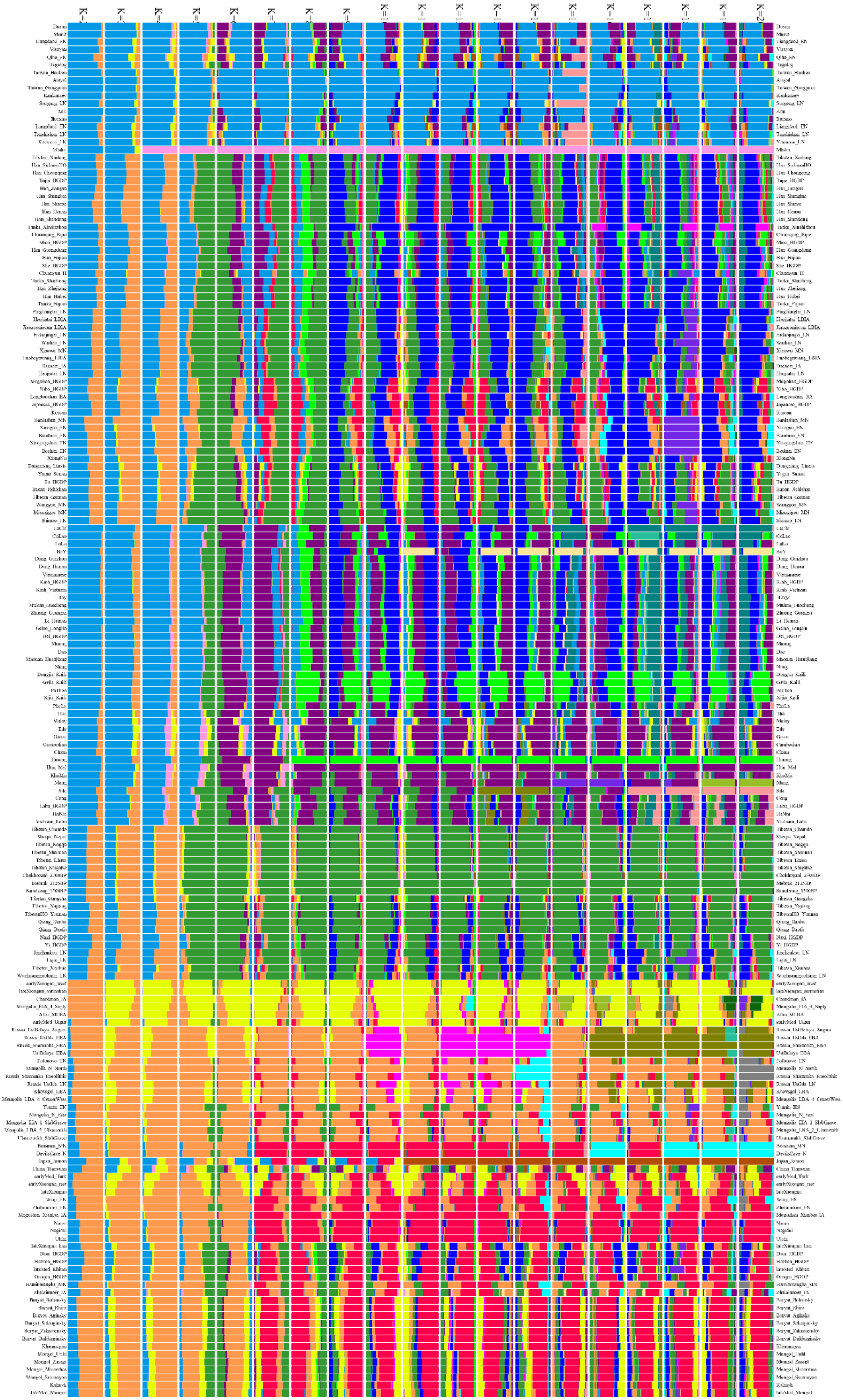

**Figure S3. Results of model-based ADMIXTURE analyses results with the predefined ancestral sources ranging from two to twenty.**

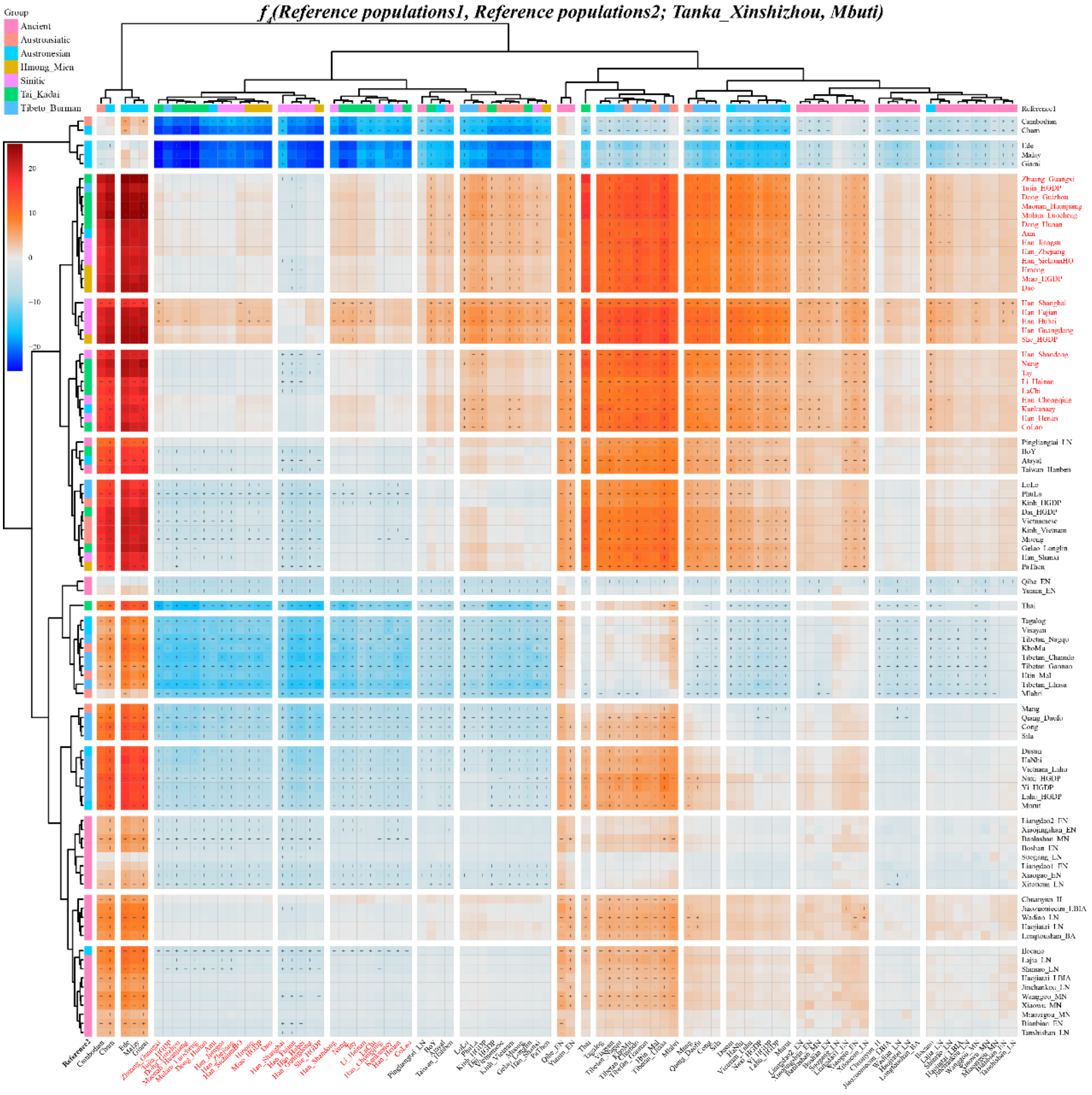


**Figure S4. Formal test of genomic affinity in Xinshizhou Tanka people inferred from the two-population comparison $f_4$-statistics in the form *$f_4$(Reference population1, Reference population2; Tanka_Xinshizhou, Mbuti)*.** Red color denoted the positive $f_4$-values, which suggested Xinshizhou Tanka people shared more derived mutations with reference population1 (left population lists), and blue color showed the negative $f_4$-values, which suggested Xinshizhou Tanka people shared more alleles with reference population2 (bottom population lists). Statistically significant results were marked with the '+'.

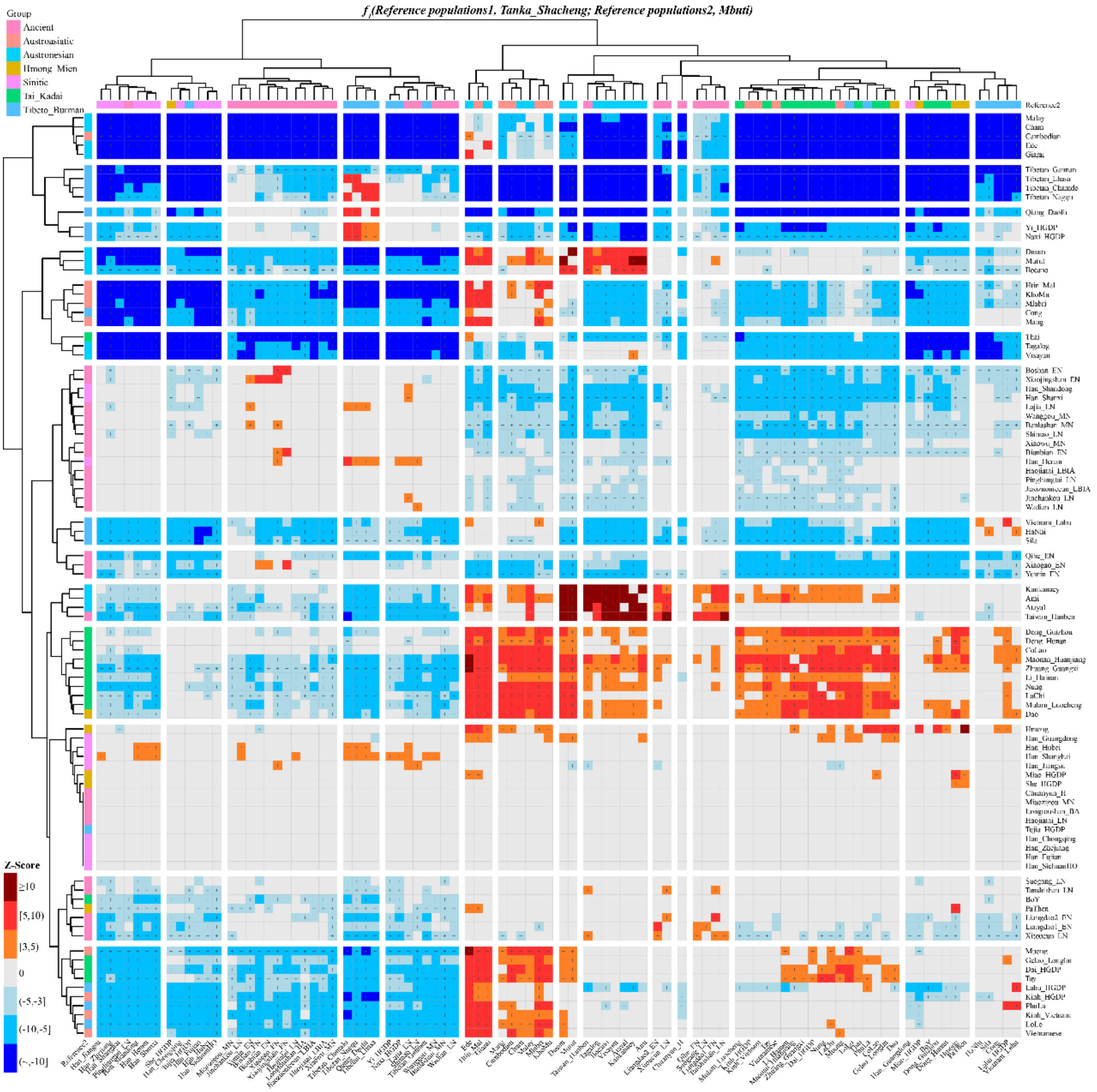

**Figure S5. A formal test of genomic continuity and admixture in Shacheng Tanka people inferred from the two-population comparison $f_4$-statistics in the form $f_4$(Reference population1, Reference population2; Tanka_Shacheng, Mbuti).** Red color denoted the positive $f_4$-values, which suggested reference population2 (bottom population lists) shared more derived mutations with reference population1 (left population lists), and blue color showed the negative $f_4$-values, which suggested reference population2 shared more alleles with Shacheng population and gray color showed no statistically significant results were observed. Statistically significant results were marked with the '+'.

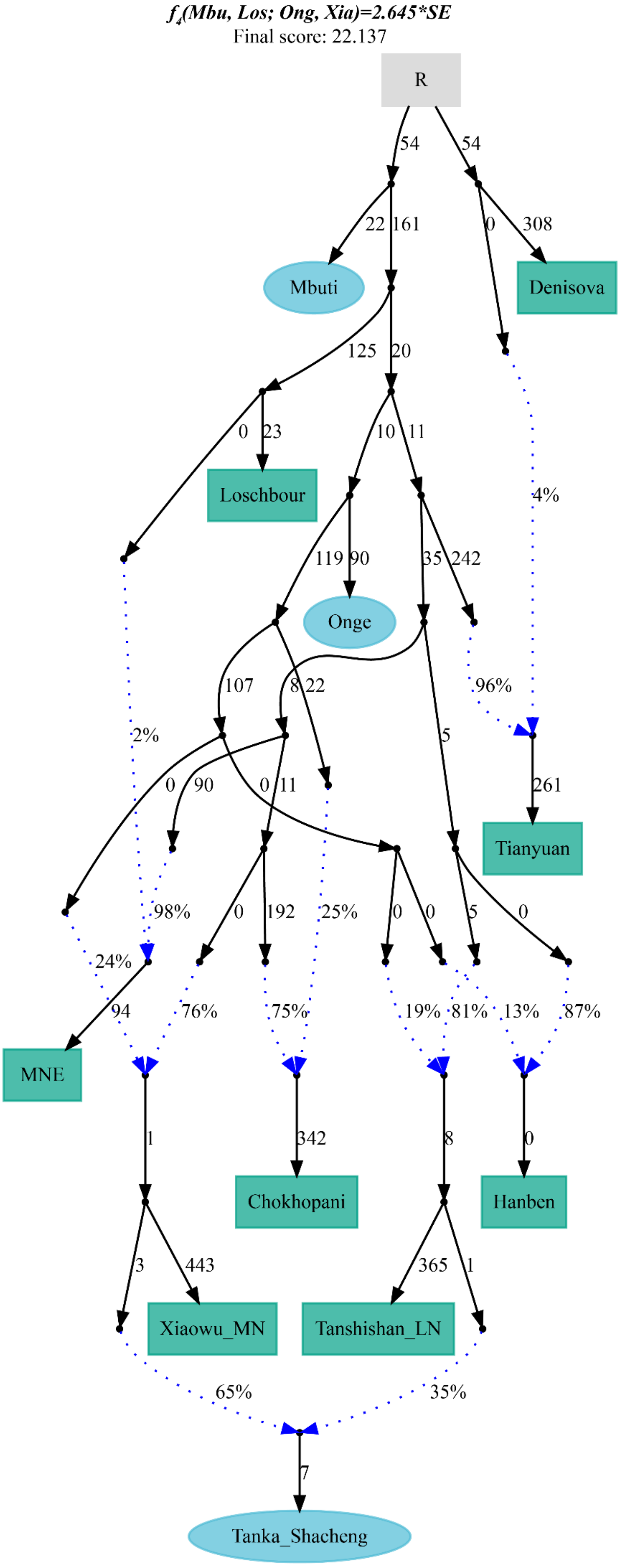

f4(Mbu, Los; Ong, Xia)=2.645*SE
Final score: 22.137
R
Mbuti
Denisova
Loschbour
Onge
Tianyuan
MNE
Chokhopani
Hanben
Xiaowu_MN
Tanshishan_LN
Tanka_Shacheng
4%
96%
2%
98%
24%
76%
75%
25%
19%
81%
13%
87%
65%
35%

**Figure S6. Genetic drift-based phylogenetic phylogeny showed population split and gene flow events for Shacheng Tanka.** Tanka people were modeled as the admixture of two Neolithic East Asian lineages. Genetic drift was marked as 1000 times of $f_2$ values. Dot blue lines denoted the admixture events and corresponding admixture proportions were also marked.
